# Discovery of a chemosynthetic community at the serpentinite-hosted seep on Quaker Seamount, Mariana Forearc

**DOI:** 10.64898/2025.12.26.696545

**Authors:** Chong Chen, Hiromi Kayama Watanabe, Jade Castel, Didier Jollivet, François H. Lallier, Margaux Mathieu-Resuge, Loïc N. Michel, Yusuke Motomura, Seitaro Ono, Junichi Miyazaki, Ken Takai

**Affiliations:** X-STAR, Japan Agency for Marine-Earth Science and Technology (JAMSTEC), 2-15 Natsushima-cho, Yokosuka, Kanagawa 237-0061, Japan; Sorbonne Université, CNRS, Station Biologique, 29680 Roscoff, France; Univ Brest, Ifremer, BEEP, F-29280 Plouzané, France; Animal Systematics and Diversity, Freshwater, and Oceanic Sciences Unit of reSearch (FOCUS), University of Liège, Liège, Belgium; Atmosphere and Ocean Research Institute, The University of Tokyo, 5-1-5 Kashiwanoha, Kashiwa, Chiba 277-8564, Japan; Department of Natural Environmental Studies, Graduate School of Frontier Sciences, The University of Tokyo, 5-1-5 Kashiwanoha, Kashiwa, Chiba 277-8563, Japan; Geological Survey of Japan, National Instutute of Advanced Industrial Science and Technology, 1-1-1 Higashi, Tsukuba, 305-8561, Japan

**Keywords:** biodiversity, chemosynthesis-based ecosystem, chemosynthetic, community composition, deep sea, hydrocarbon seep, mud volcano, serpentinisation, Western Pacific

## Abstract

Deep-sea chemosynthesis-based ecosystems in the Mariana Region include hydrothermal vents on the Mariana Arc and in the Mariana Back-Arc, and serpentinite-hosted seeps on the Mariana Forearc – with the latter being by far the least studied. Here, we surveyed the biodiversity of the serpentine seep on the Quaker Seamount. At first glance, the area appears to be devoid of megafauna, but examination of the carbonates revealed dense aggregations of animals dominated by the limpets *Bathyacmaea* and *Pyropelta* on brucite-carbonate chimney structures exhibiting visible fluid venting. Sorting of recovered material yielded a total of 14 species and together with an observation of a *Munidopsis* squat lobster we report a total of 15 species; six of these are considered endemics of chemosynthetic ecosystems. The occurrence of taxa such as the snail *Lurifax* cf. *japonicus* and the limpet *Pyropelta ryukyuensis*, otherwise only known from distant vents in the Izu-Ogasawara Arc and Okinawa Trough off Japan, strengthens the hypothesis that serpentine seeps may function as dispersal stepping-stones across arc and back-arc systems.

## Introduction

Chemosynthesis-based ecosystems in the deep sea such as hydrothermal vents and hydrocarbon seeps support a unique and largely endemic fauna that do not occur in the ambient seafloor (Tunnicliffe et al. 1998; Levin 2005). These systems are usually associated with geological activities and powered by the discharge of geofluids containing reduced compounds such as hydrogen sulfide and methane, which microbes oxidise to obtain chemical energy for carbon fixation (Sogin et al. 2021). Animals living in chemosynthetic habitats often form metapopulations across multiple sites connected by larval dispersal, and sites from different biogeographic regions around the world host distinct suites of species (Rogers et al. 2012; Brunner et al. 2022; Tunnicliffe et al. 2023).

The Mariana Trench-Arc system represents a non-accretionary convergent plate margin where the Pacific Plate subducts beneath the Philippine Sea Plate, and is among the world’s most tectonically dynamic locations (Ohara et al. 2012). The system can be divided into several features with distinct underlying geology, with Mariana Trench at the eastern end and followed by Mariana Forearc, Mariana Volcanic Arc, Mariana Back-arc (Trough) spreading centre, and the West Mariana Ridge remnant arc towards the west (Embley et al. 2007). Both the volcanic arc and the back-arc host numerous active hydrothermal vent communities, but only few animal species are able to live in both settings and most appear to be restricted to either one (Giguère and Tunnicliffe 2021).

The forearc, on the other hand, is home to several serpentinite-hosted cold seeps with chemosynthesis-based animal communities, including the 5700 m deep Shinkai Seep Field on the southern forearc (Ohara et al. 2012) and two large serpentinite mud volcanoes on the northern forearc including the 2900 m deep South Chamorro Seamount (Fryer and Mottl 1997; Chen et al. 2024c) and the 1200 m deep Asùt Tesoru Seamount (Chen et al. 2023). These serpentinite-hosted systems are characterised by strongly alkaline geofluids with pH of up to 12.5 and rich in hydrogen gas and methane (Mottl et al. 2003; Okumura et al. 2016; Fryer et al. 2018; Eickenbusch et al. 2019). The three sites differ greatly in species composition, hypothesised to reflect the depth differences more than the distances between the sites (Chen et al. 2024c). Some species found at these serpentinite-hosted seeps have affinities to both much more northern locations (e.g., the vesicomyid clam *Ectenagena nautilei* first described from Nankai Trough seeps off Japan) (Chen et al. 2024c) and more southern locations (e.g., the provannid snail *Provanna buccinoides* previously only known from the southern Pacific hydrothermal vents) (Chen et al. 2023), indicating the potential role of these seeps as dispersal ‘stepping-stones’ between biogeographic regions.

Locating and surveying more serpentinite-hosted seep faunal communities on the Mariana Forearc is key in assessing the role of these seeps in the biogeography of chemosynthetic ecosystems across the entire western Pacific. There are many more serpentinite mud volcanoes on the northern Mariana Forearc (Fryer 2012), but they have not been targeted by submersible dives focusing on the biology and it remains unclear whether they host chemosynthetic faunal communities. On active serpentinite-hosted seeps the geofluid seepage often leads to the formation of carbonate and brucite chimneys (Kelley et al. 2007), and such chimneys have been observed on other Mariana serpentinite mud volcanoes including Conical Seamount, Pacman Seamount (the Cerulean Seep), and Quaker Seamount (Fryer et al. 1990; Fryer et al. 1999; Fryer 2012). These seamounts are thus likely actively seeping, making them potential candidates for hosting seep animals.

Here, we report and characterise the fauna endemic to chemosynthetic ecosystems in the serpentinite-hosted cold seep on the Quaker Seamount. This seamount was previously visited by two dives using the remotely operated vehicle (ROV) *Jason II* (Fryer et al. 2006; Tong et al. 2021), with brief reports on the chimney mineralogy but there was no indication of the presence of chemosynthetic fauna. We further discuss the biogeographic implications of the fauna at Quaker, and how it compares with the three other known serpentinite-hosted seeps of the Mariana region.

## Materials and Methods

Quaker Seamount was explored during the cruise KM25-12 “MoWAME” aboard the research vessel (R/V) *KAIMEI* which took place between December 2025 to January 2026. Seafloor bathymetry was collected with a Kongsberg EM122 multibeam echosounder installed on the vessel, and the data were subsequently cleaned by CARIS HIPS and SIPS version 11.4 to produce the final bathymetric map at 10 m grid resolution using the software GMT6 (Wessel et al. 2019). A single ROV dive (#357; December 24, 2025) using the *KM-ROV* was carried out at Quaker Seamount, during which seafloor footage was recorded with a high-definition (1920×1080 pixels) camera (Mini Zeus: Mini Colour HD Camera) and an 8K (7680×4320 pixels) 180° fisheye video camera (Z CAM E2-F8). As the *KM-ROV* lacked a still camera, seafloor images were extracted as frame grabs from the video recording. In addition, a stand-alone RINKO II dissolved oxygen (DO) and temperature sensor (JFE Advantech Co., Ltd.) was used to take basic environmental parameters of the animal aggregations; measurements were taken over a two-minute period with data recorded every second. The ROV dive track was then mapped on our new bathymetry.

Chemosynthesis-based faunal assemblages were sampled by the *KM-ROV* either by a suction sampler or by directly collecting chimneys to which animals were attached using a manipulator arm, and placing them in a sample box. Upon recovery on-board, the animals were immediately moved into a cold room set to 4°C. The samples, including seawater in the boxes, were sieved on a 0.5 mm sieve to retrieve macrofauna samples. As we focused on macrofauna in this study, it is probable that meiofauna taxa were present but are not accounted for here. The sieved material was sorted and identified with the aid of a dissecting microscope (Olympus SZX10), and then preserved in 99% ethanol, 10% buffered formalin, or frozen in -80°C deep-freezer. Photographs of larger animals were taken using a Canon EOS-5Ds R digital single-lens reflex camera mounted with a macro lens (Canon EF 100 mm F2.8L MACRO IS USM), and those of smaller ones were captured by a CMOS camera (Advan Vision AdvanCam-HD2sA) attached on the trinocular of the dissecting microscope.

## Results and Discussion

### Dive survey at Quaker Seamount

We revisited the summit of Quaker Seamount (Figure 1A-B) previously dived by ROV *Jason II* in 2003 (Tong et al. 2021), where a brucite-carbonate mound topped by dense clusters of irregular, finger-like columnal spires was reported (Fryer 2012). We were able to locate this mound (Figure 2A-B), which we named the Miclas chimney (18°45.3479’N, 146°59.0003’E, 2216 m deep; Figure 1B). Our observations revealed visible venting of shimmering geofluid from the tip of the spires (Figure 2E), clearly showing this site exhibits active fluid discharge.

**Figure 1.**
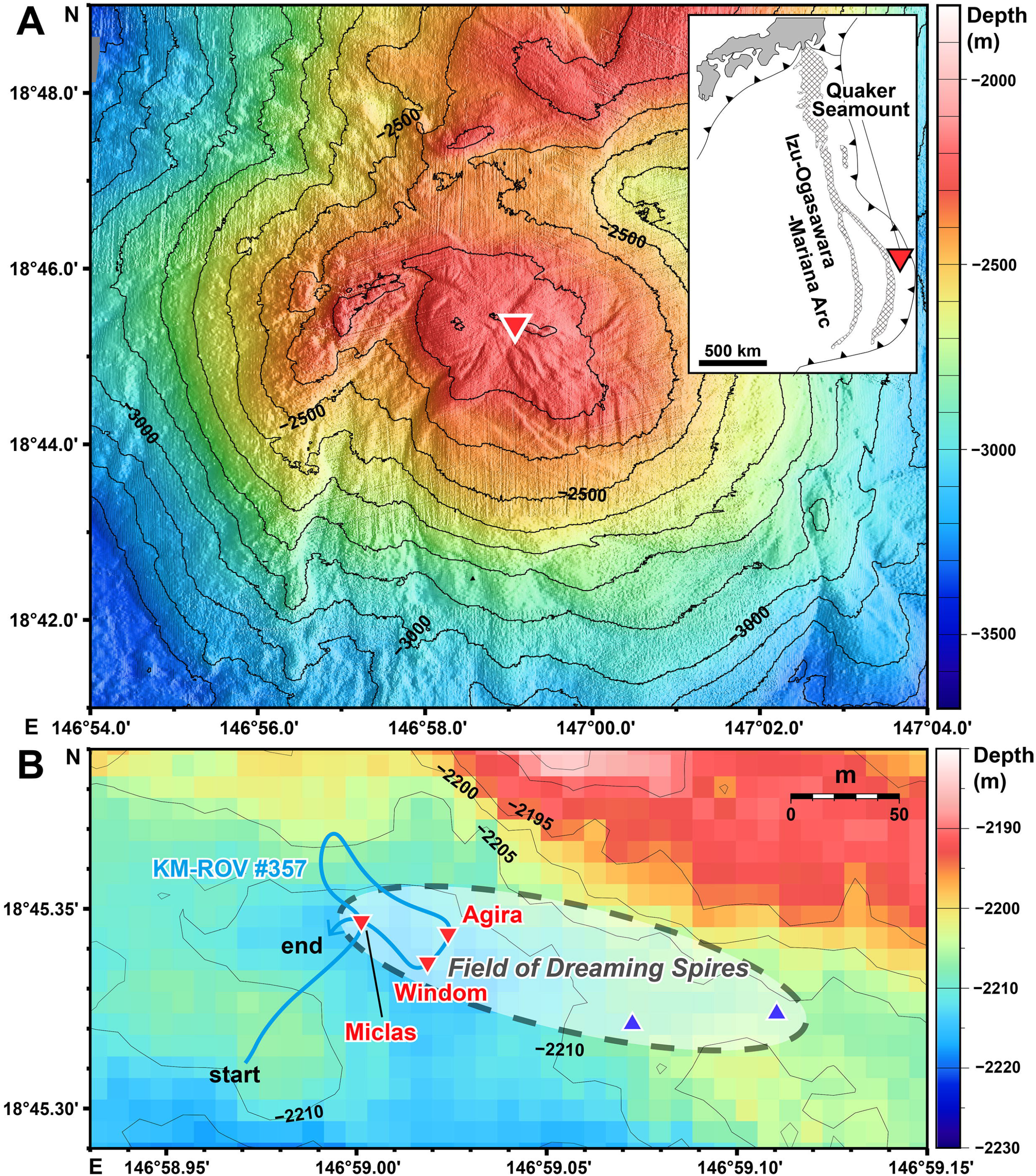
Bathymetric map generated from shipboard multibeam data of R/V *Kaimei* showing **A**) Quaker Seamount indicating the location of the Field of Dreaming Spires (red inverted triangle); and **B**) the extent of the Field of Dreaming Spires (grey dotted line), showing the dive track of *KM-ROV* dive #357 (blue line), the three main chimney structures (red inverted triangles), and the two small chimneys found during previous surveys of ROV *Jason II* in 2003 (blue triangles).

**Figure 2.**
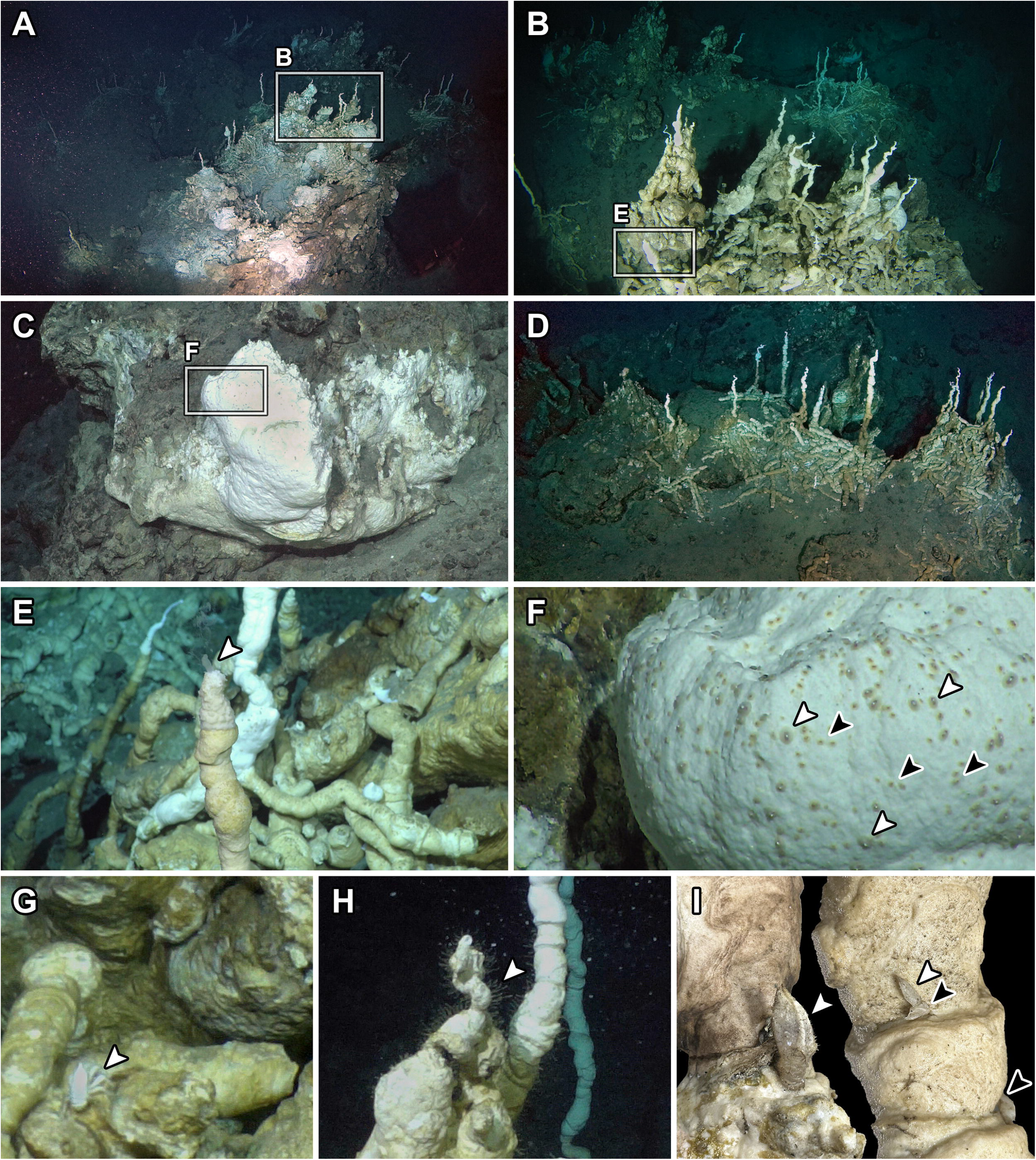
*In situ* observations of the Field of Dreaming Spires serpentinite-hosted seep on Quaker Seamount. **A)** Overview of the Miclas chimney showing finger-like spires on and around the main mound; **B)** a close-up showing the top of the Miclas chimney. **C)** The Agira chimney. **D)** The Windom chimney. **E)** A spire on the Miclas chimney exhibiting visible active venting of shimmering geofluid (white arrowhead). **F)** A dense aggregation of limpets (white arrowheads: *Bathyacmaea* sp. ‘South Chamorro’ *sensu* (Sato et al. 2020); black arrowheads: *Pyropelta ryukyuensis*) on the surface of the Agira chimney. **G)** A squat lobster in the genus *Munidopsis* spotted on the Miclas chimney. **H)** Dense coverage of hydroids (Leptothecata indet.) on spires of the Miclas Chimney. Surface of spires collected from the Windom chimney showing the barnacles *Trianguloscalpellum* sp. (white arrowheads) and *Altiverruca* sp. (black arrowheads).

Previous analyses indicated younger spires at Quaker are dominated by calcite and magnesium-rich alkaline minerals like brucite and hydromagnesite, being enriched in magnesium oxides (18.5-37.5%) but depleted in calcium oxides (12.2-32.1%) (Tong et al. 2021). As chimneys mature, they show lower magnesium oxide contents (1.5-23.6%) and higher calcium oxide contents (18.6-53.3%), coinciding with increased aragonite in the mineral composition (Tong et al. 2021). The inner core of the distal tip is thought to be the most recently formed part comprising mostly of magnesium-rich minerals, which agrees with our observations.

Although visible fluid discharge is known from other serpentinite-hosted hydrothermal systems such as the Lost City field on the Mid-Atlantic Ridge (Kelley et al. 2007) and the Kunlun hydrothermal system near the Mussau Trench (Li et al. 2025), this is the first known case from the Mariana Forearc (Okumura et al. 2016) and marks Quaker as a remarkable target for future geochemistry and microbiology research. Previously, pH of 9.2 was recorded from piston core porewater samples taken at Quaker (Hulme et al. 2010), and pH of pure discharges from these chimneys could be even higher.

On the Miclas chimney, the dominant fauna were two species of limpets whose genera are endemic to chemosynthetic systems – *Bathyacmaea* sp. ‘South Chamorro’ *sensu* Sato et al. (2020) and *Pyropelta ryukyuensis* (Chen et al. 2025) (Figures 2F, 3A-B). Both temperature (1.89°C, standard deviation (SD) = 0.0010) and DO (4.2036 mg/L, SD = 0.0014) at the limpet aggregation did not differ significantly from the reference measurements away from active seepage (1.88°C, SD = 0.0010; 4.2059 mg/L, SD = 0.0015). Nevertheless, we note that the minimum distance between the sensor and the chimney surface we can realistically achieve is about 3 mm, and it is possible that the microchemical conditions experienced by the limpets are different. We also spotted a squat lobster in the genus *Munidopsis* (Figure 2G) and numerous hydroids (Leptothecata indet.) covering the chimneys (Figure 2H).

**Figure 3.**
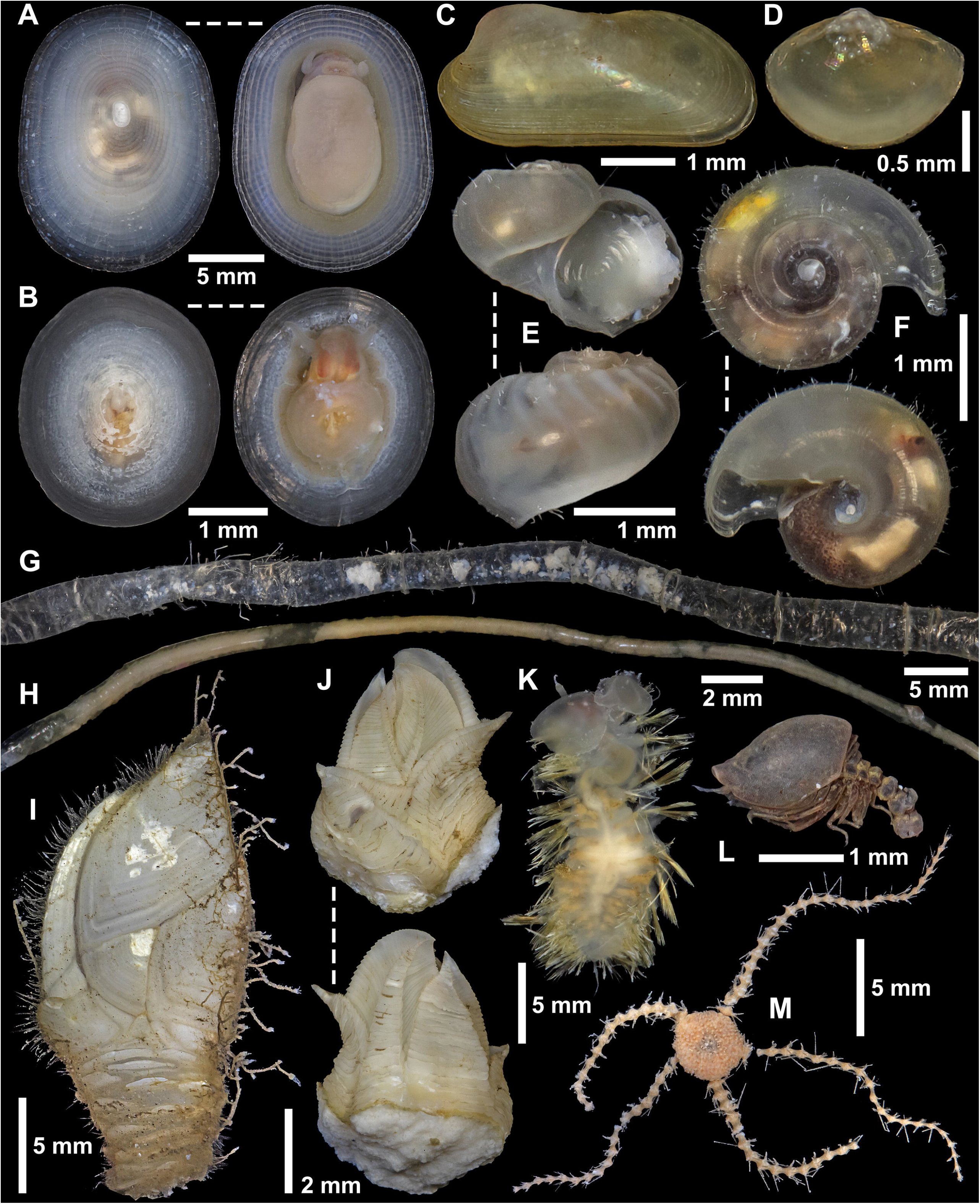
Animals collected from the Field of Dreaming Spires seep on Quaker Seamount. **A)** *Bathyacmaea* sp. ‘South Chamorro’ *sensu* (Sato et al. 2020); **B)** *Pyropelta ryukyuensis*; **C)** *Gigantidas* sp. juveniles; **D)** the protobranch bivalve *Yoldiella* sp.; **E)** Neomphalidae indet.; **F)** *Lurifax* cf. *japonicus*; **G)** the parchment worm *Spiochaetopterus* sp.; **H)** the siboglinid tubeworm *Siboglinum*? sp.; **I)** the stalked barnacle *Trianguloscalpellum* sp. with hydroids (Leptothecata indet.) growing on it; **J)** the barnacle *Altiverruca* sp.; **K)** Polynoidae indet.; **L)** Cumacea indet.; **M)** Ophiuroidea indet.

Approximately 40 m east of Miclas (Figure 1B), we encountered a different type of carbonate chimney where a large block of white carbonate material was formed instead of numerous finger-like spires (Figure 3C), similar to those seen in the Shinkai Seep Field (Okumura et al. 2016). We called this the Agira chimney (18°45.3407’N, 146°59.0143’E, 2213 m deep). The surface of the Agira chimney was notable in hosting a very dense aggregation of the same two limpets as seen on Miclas. On this aggregation the temperature (1.87°C, SD = 0.0020) and DO (4.1980 mg/L, SD = 0.0013) also did not differ notably from the reference measurement.

Going southwest for about 20 m from Agira revealed another chimney complex of numerous finger-like spiral chimneys similar to those seen on and around Miclas (Figure 2D). We named this site the Windom chimney (18°45.3359’N, 146°59.0081’E, 2218 m deep) and proceeded to sample chimneys, however this site did not exhibit any dense aggregations of limpets seen on the other two chimney sites. On chimneys from this site, we found two species of barnacles including *Trianguloscalpellum* sp. (Figure 3I) and *Altiverruca* sp. (Figure 3J). Although they were attached to the carbonate-brucite chimneys which were actively seeping (Figure 2I), these are both genera that are common in the ambient seafloor and are not restricted to chemosynthesis-based ecosystems (Chan et al. 2010). They are probably filter-feeding opportunists that utilise the increased food availability and availability of hard substrates at the seep site.

As we discovered a faunal community characteristic of chemosynthesis-based ecosystems, we name this seep site the Field of Dreaming Spires. Due to the ROV dive time restrictions we were unable to transit further east to revisit the two locations where small chimneys were seen in 2003 (Figure 1B). Nevertheless, from these previous observations we deduce that the seepage activity on this site spreads over about a 200 m line from west to east.

### Species Composition

Sorting the material collected on-board the research vessel revealed further species that co-occurred with the two dominant limpets. In total, we sampled 14 animal species from the Field of Dreaming Spires (Figure 3). Together with the *Munidopsis* squat lobster that was not collected, we counted a total of 15 species from this site. Of these, there were two gastropods, including *Lurifax* cf. *japonicus* (Figure 3F) and Neomphalidae indet. (Figure 3E). *Lurifax japonicus* is a species so far only known from the Sumisu Caldera in the Izu-Ogasawara Arc off Japan over 1300 km to the north of Quaker Seamount (Chen et al. 2024a), and the specimens from Quaker are very similar in terms of shell morphology although they could be genetically divergent. As this genus is not known from either Mariana Arc or Back-Arc hydrothermal vents (Giguère and Tunnicliffe 2021), it is a surprise to find it on a serpentinite-seep on the Forearc. The neomphalid snail is morphologically identical to the one recovered from Asùt Tesoru Seamount, another serpentinite seamount on the Mariana Forearc (Chen et al. 2023) and likely belongs to the same species in an undescribed genus. Another similar species was collected from the Japan Trench, where the presence of a seep site has been hypothesised (Fukumori et al. 2019) – this may be a seep-specialist genus.

A few specimens of a bathymodioline mussel in the genus *Gigantidas* (Figure 3C) were found attached to spiral chimneys at Miclas. All were small juveniles between 2-5 mm in shell length, and we did not observe any adult individuals during the ROV survey. As virtually all bathymodioline mussels rely on endosymbiotic bacteria for energy (Lorion et al. 2013), it may be that this species is capable of dispersing and settling at Quaker but there is no sufficient hydrogen sulfide input for the mussels to grow to adulthood. An undescribed *Gigantidas* species inhabits the more southern South Chamorro seamount serpentinite-seep (Yamanaka et al. 2003a; Yamanaka et al. 2003b; Chen et al. 2024c), but the Quaker specimens were too small for accurate identification to the species level. Another bivalve collected was a protobranch in the genus *Yoldiella* (Figure 3D).

Three annelid species were sampled. This included the chaetopterid parchment worm *Spiochaetopterus* sp.; chaetopterid worms like *Phyllochaetopterus* cf. *polus* (Figure 3G) have been found at many sites influenced by serpentinization such as the Ashadze vent field on the Mid-Atlantic Ridge and the Shinkai Seep Field (Morineaux et al. 2010; Watanabe et al. 2021). We only collected one living specimen each of the small siboglinid worm *Siboglinum*? sp. (Figure 3H) and an unidentified polynoid worm recovered in damaged condition (Figure 3K). *Siboglinum* was previously also found on the Asùt Tesoru Seamout (Chen et al. 2023), while polynoids are common members of chemosynthetic communities including serpentinization seeps like Asùt Tesoru and South Chamorro Seamounts (Chen et al. 2024c).

Finally, we also collected three species which we cannot be sure whether they are specific to chemosynthetic ecosystems or not, including a hydroid (Figure 3I, living on the barnacle *Trianguloscalpellum*), a cumacean (Figure 3L), and a brittle star (Figure 3M). Overall, at least six species are likely specific to chemosynthetic ecosystems (*Bathyacmaea* cf. “South Chamorro” *sensu* Sato et al. (2020), *Pyropelta ryukyuensis, Gigantidas* sp., Neomphalidae indet., *Lurifax* cf. *japonicus, Siboglinum*? sp.). The complete lack of abyssochrysoidean snails such as *Provanna* and *Desbruyeresia* is notable, as they are common in other serpentine mud volcanoes.

### Biogeographic Implications

In the context of the three previously characterised forearc sites including Asùt Tesoru (∼1200 m), South Chamorro (∼2900 m), and the much deeper Shinkai Seep Field (∼5700 m), Quaker (∼2200 m) helps bridge the large depth gap that has been hypothesised to structure the faunal community (Chen et al. 2024c). Table 1 presents the presence and absence of major taxonomic groups linked to chemosynthetic habitats found at these serpetinite-hosted seeps. The Quaker assemblage is most similar to Asùt Tesoru, sharing multiple seep-affiliated taxa (including *Pyropelta, Bathyacmaea*, Neomphalidae indet., and *Siboglinum*), consistent with a depth-filtered serpentinite-hosted seep fauna. South Chamorro Seamount hosts the most diverse taxa especially more chemosymbiotic bivalves, which is likely linked to the higher fluid discharge at this site compared to Quaker or Asùt Tesoru; and likely only a limited number of taxa is capable of thriving at the 5700 m depth of Shinkai Seep Field. The repeated occurrence of chaetopterid parchment worms at these seeps is of interest and they may have a preference for serpentinite-hosted sites. While these dominant taxa are mostly shared with non-serpentinite seeps at the genus level, genera that characterise inactive hydrothermal vents such as *Neolepetopsis* limpets and *Melanodrymia* snails (Chen et al. 2024b; Mullineaux et al. 2025) are lacking at serpentinite seeps.

**Table 1.**
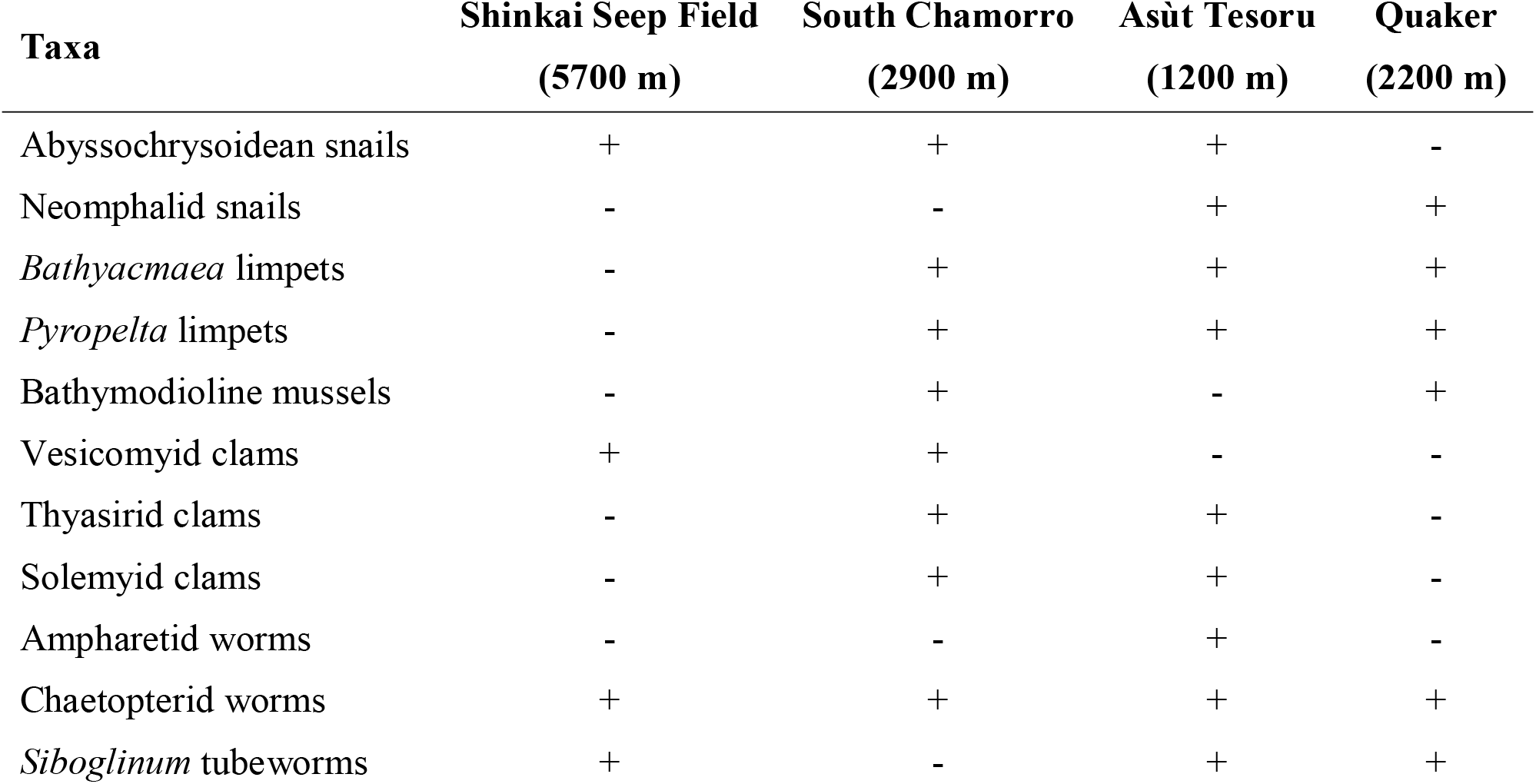
Presence-absence of main animal groups linked to chemosynthetic habitats at Quaker Seamount and other known Pacific serpentinite-hosted seeps (Onishi et al. 2018; Chen et al. 2023; Chen et al. 2024c).

At the same time, several taxa at Quaker suggest that serpentinite-hosted seeps may connect chemosynthetic communities in the western Pacific (Tunnicliffe et al. 2023) in unexpected ways. The occurrence of *Lurifax* otherwise only known from Sumisu Caldera in the Izu–Ogasawara Arc among the nearby sites in the northwestern Pacific, as well as another instance of finding *Pyropelta ryukyuensis* in the Mariana Forearc seeps outside of the Okinawa Trough vents (Chen et al. 2025), provide counter-examples against the view that western Pacific vent fauna are largely restricted to their biogeographic features (e.g., arc, back-arc, basin) (Mitarai et al. 2016) and that arc and back-arc fauna rarely mix (Giguère and Tunnicliffe 2021; Tunnicliffe et al. 2023). These lend support to the hypothesis that serpentinization-hosted seeps may be important dispersal stepping stones for hydrothermal fauna and vice versa (Chen et al. 2024c), while why some species capable of living at vents occur at these seeps but not at other Mariana vents requires further research. The presence of *Provanna buccinoides* in the South Chamorro serpentine seep, otherwise only known from southern Pacific vents, further supports this pattern (Chen et al. 2024c). Additional faunal survey at other active serpentinite mud volcanoes in the Mariana Forearc such as the Cerulean Springs seep on the Pacman Seamount (Fryer et al. 1999; Fryer 2012), will further clarify the roles of serpentinite-hosted seeps in the biogeography of chemosynthetic ecosystems.

## Acknowledgements

We are grateful to the captain and crews of R/V *KAIMEI* during the research cruise KM25-12 “MoWAME”, and extend our thanks to the pilots and technical teams of ROV *KM-ROV* for their support of our sample collection. The JAMSTEC outreach team for the cruise (Natsumi Nakano, Rena Murata, and Yoshiyuki Ogita) are acknowledged for their great help in preparing and deploying the 8K video camera (Z-Cam). We thank Eijiroh Nishi (Yokohama National University) for help in identifying the chaetopterid worm; constructive comments from Lisa Levin (Scripps Institution of Oceanography) and an anonymous reviewer helped improve an earlier version of this paper. We take this opportunity to pay tribute to the Japanese TV series *Ultraseven* (Tsuburaya Productions, 1967-1968), for captivating our imagination and introducing the idea of Capsule Kaiju which we borrowed the names of the chimneys from.

## Author Contributions

CC conceived and designed the project. CC, JM, and KT participated in dive planning and leading the dive survey. CC and HKW, JC, DJ, FL, MM-R, LNM, and YM collected, sorted, identified, and interpreted animals from the cold seep community. JC, DJ, FHL, MM-R, LNM, and YM helped in sorting the specimens. SO analysed multibeam data and generated the bathymetric map. SO and CC interpreted and mapped the dive tracks. KT supervised the project and led the research cruise. CC drafted the original manuscript which was critically edited by all other authors. All authors agreed to the submission and publication of the present manuscript.

## Statements and Declarations

### Competing Interests

We have no competing interests to declare.

### Ethical Statement

The study species are invertebrate animals and no experimental manipulation was undertaken on live animals in this study. All applicable international, national, and/or institutional guidelines for the care and use of animals were followed by the authors. Sampling on Quaker Seamount, within the Commonwealth of the Northern Mariana Islands, was carried out under the Marine Scientific Research permit number MSR U2025-017 from the United States government.

### Data Availability

All data generated or analysed during this study are included in this published article or in the submersible dive video footage, which is publicly available on YouTube: https://www.youtube.com/watch?v=EMwpv2qbt_w.

## Notes

https://www.youtube.com/watch?v=EMwpv2qbt_w

### Competing Interest Statement

The authors have declared no competing interest.

### Summary of Updates

Revised version submitted to the journal Marine Biodiversity.

